# Vocally mediated coordination during a cooperative task in parrots

**DOI:** 10.1101/2025.10.10.681640

**Authors:** Sara Torres Ortiz, Thorsten Johannes Skovbjerg Balsby, Wesley Webb, Matthew Pawley, Ole Næsbye Larsen

## Abstract

Studying vocal coordination in a cooperative setting provides insights into how animals process social information, coordinate their actions, and make decisions in complex social settings. It can also reveal how animals use vocal signals to navigate complex social interactions that might not be apparent in solitary or non-cooperative contexts. To our knowledge, experimental evidence for intentional vocal communication in controlled experiments has been demonstrated only in dolphins. Here, we investigate whether peach-fronted conures (*Eupsittula aurea*) can use vocal communication to coordinate behaviour in a cooperative task. We chose parrots due to their complex social systems, their vocal learning and mimicry skills, and their advanced cognitive abilities. We tested four individuals pairwise in all combinations in the loose string paradigm, which requires synchronization of the two individuals to retrieve a food reward by simultaneously pulling a string. To verify that the parrots used vocal communication to solve the task, we tested them first under three control conditions where visual information and simple behaviour sufficed to solve the tasks. We then compared their vocal behaviour in the control conditions with their vocal behaviour in a task without visual information. During performance of the latter task, the birds demonstrated significantly higher call rates and variability and specific associations between call types and trial outcome compared to the control conditions. After the performance of the task, the birds used vocal convergence more often after failed trials, potentially as a reconciliation mechanism. Our results indicate that peach-fronted conures can use vocalizations to coordinate the solving of a cooperative task, and that vocal convergence may serve as a bonding mechanism following unsuccessful cooperative efforts.

## Introduction

Communication is a social event where an emitter produces a signal to be received by a listener with the purpose of influencing the listener’s behaviour; communication has occurred when the listener receives and responds to the emitted signal (Seyfarth & Cheney, 2003). The meaning and function of those signals may differ from the perspective of the emitter and the listener (Marler, 1961). To assess the possible intention of the signaller and reaction of the listener, communication in non-human animals is often studied in behavioural contexts, where the general behaviour is linked with the vocalization type (Henderson et al., 2011). Using this approach, researchers have been able to identify many different types of vocalizations such as mating and alarm calls. In these cases, the meaning and function of the calls is clear, however, a different methodology is needed to understand the call usage in more general situations. For example, when socializing, animals can display a great variety of vocalizations and even though it is possible to identify the emitter, it is not always clear what the meaning and function of the vocalization is, or for whom the call is intended.

It has been hypothesised that cooperation and communication coevolved in humans, where it is often argued that communication appeared in order to facilitate coordination of behaviour (Hayes & Sanford, 2014; Miller et al., 2002; Salahshour, 2020). It has been hypothesized that advances in human communication facilitated development of more complex cooperative behaviour, which in turn selected for more sophisticated linguistic skills (Hayes & Sanford, 2014; Tomasello et al., 2012). In many animal species, vocalizations play a crucial role in coordinating group behaviours. For example, African wild dogs (Walker et al., 2017), meerkats (Bousquet et al., 2011), red-fronted lemurs (Sperber et al., 2017), and jackdaws (Dibnah et al., 2022) use vocal signals to facilitate consensus and trigger group movements or other collective actions (Chereskin et al., 2024). In some cases, vocalizations not only help coordinate group activities but also enable cooperation to accomplish shared tasks to achieve common goals (King et al., 2021). In non-human animals, so far, vocal communication during a cooperative task has been studied in only one species: bottlenose dolphins (*Tursiops truncatus*). Using a cooperative task where the individuals had to synchronize their actions to push two buttons at the same time to obtain a reward, vocal communication in dolphin pairs was analyzed (Eskelinen et al., 2016; King et al., 2021). The number of whistles produced by the dolphins increased significantly right before pressing the button suggesting that vocal communication was used for behavioral synchronization. However, differences in communication and performance between trials with and without visual feedback were not analyzed. This means that the dolphins could have used the whistles, for instance, as a reinforcement but relied on visual feedback or other cues for coordination.

In complex societies such as those of anthropoid primates, strong social bonds are vital (Wittig et al., 2007). After conflict, relationships are repaired through a process called “reconciliation”, involving grooming, touching, or vocalizing (Cheney & Seyfarth, 1989; Silk, 2002; Wittig & Boesch, 2005). Whether birds use similar mechanisms remains unclear, but some studies indicate similar reconciliation strategies in socially complex bird species, such as corvids and tits (Sima et al., 2018; Williams, 2024). For example, ravens (*Corvus corax*) respond vocally in different ways to group members, which they have not encountered for up to three years compared to strangers, with their response depending on the nature of their prior relationship (Boeckle & Bugnyar, 2012). Reconciliatory behaviors, such as contact sitting, preening or beak-to-beak or beak-to-body touching, commonly observed among ravens with strong social bonds, further highlight the role of these behaviors in mending damaged relationships (Fraser & Bugnyar, 2011).

Parrots show many cognitive abilities similar to those observed in large-brained mammals such apes and dolphins (Pepperberg, 1990, 2002). Regarding their vocal behaviour, they are one of three bird taxa (songbirds, parrots and hummingbirds) capable of vocal production learning, meaning that they are capable of modifying their vocalizations, for instance, by imitating others. Parrots have been observed producing sequences of calls with patterns of increasing or decreasing similarity to the calls of the vocal partner. The systematic convergence or divergence of acoustic features may carry different meanings for these parrots, and it has been proposed that such rapid vocal modifications could help mediate short-term affiliations and may play a role in negotiating dominance or group decisions (Balsby & Scarl, 2008; Bradbury & Balsby, 2016). Like songbirds with open-ended song learning parrots have the ability to modify their vocal repertoires throughout their lives. Parrots also rely heavily on vocal communication, since their gestural repertoire is limited (Moura et al., 2014) and have diverse repertoires that include individual-specific contact calls. (Thomsen et al., 2013). Even though studies in the wild with peach-fronted conures (*Eupsittula aurea*) have not been conducted, several field studies have been conducted with orange-fronted conures (*Eupsittula canicularis*), close relatives of peach-fronted conures (e.g., Balsby & Bradbury, 2009; Balsby & Scarl, 2008; Wright et al., 2003). Their results suggest that these parrots are capable of rapid imitation of contact calls (Balsby et al., 2012; Vehrencamp et al., 2003) and that they may address a specific individual in a communication network by imitating their contact calls, like bottlenose dolphins do (King & Janik, 2013).

While parrots are an interesting animal model for investigating vocal communication, studying their vocal abilities and use in the wild is challenging. They have extensive home ranges, and forage high up in dense canopies, thus complicating visual observations. If fitted with microphones or transmitters, they are often able to remove the equipment with their powerful beaks. Therefore, laboratory studies are called for.

To gain a better understanding of the flexible use of vocal communication in parrots we recorded vocal behaviour in pairs of peach-fronted conures during a cooperative task, the loose-string task, performed under controlled conditions in the laboratory, where two birds had to coordinate their actions to solve the task and get a reward. The behavioural part of this study has been reported in Ortiz et al., 2020. Thus, the aim of the present investigation was to analyse in depth the vocalizations produced by peach-fronted conures during the task. We hypothesized that the parrots would produce more vocalizations in the test condition, where coordination is required, compared to the three control conditions where coordination is not necessary. Additionally, we expected that the type and frequency of vocalizations would differ between the control and test conditions, with the parrots using more complex or varied calls when coordination was required. Finally, we predicted that vocalization patterns would be influenced by trial success or failure, with parrots potentially using vocal imitation as a mechanism for social bonding and reconciliation after unsuccessful trials.

## Materials and Methods

### Subjects and housing conditions

Two male (M1, M2) and two female (F1, F2) peach-fronted conures were tested in pairs on the loose-string paradigm, which was developed by Hirata & Fuwa, (2007) and later used by Melis et al., (2006b, 2006a) with chimpanzees. The non-vocal behavioural results of our experiment were published in Ortiz et al. (2020). In the current investigation, we used the sound recordings obtained to test if vocal communication was used for coordination during the experiment. The parrots came from private breeders and were raised by their parents; therefore, they were not accustomed to humans and needed to be desensitized to human handling and presence. F1 and F2 hatched from the same clutch as well as M1 and M2. Data collecting experiments were conducted from November 2016 to March 2017 when the birds were between 30 for one of the pairs and 33 months old for the other pair. Before this experiment, the parrots were individually tested in the string-pulling test (Ortiz et al., 2019). The housing conditions were described in Ortiz et al. (2020).

### Setup for the behavioural experiment

The apparatus consisted of an out-of-reach sliding board with food rewards for each bird (Figure 1a, b, c). The board featured two eyelets and a wool string running through them into individual cages for each parrot, which were separated by a trap door that the experimenter could remotely operate to minimize observer bias (Sebeok & Rosenthal, 1981). The sliding board was positioned out of reach with the string available for each bird. To retrieve the food, both parrots had to pull their ends of the string simultaneously; pulling one end alone would only move the string, making the other end unavailable to the partner. All pair combinations were tested sequentially under four conditions: three control conditions (‘*Cooperative* condition’, ‘*Delayed* condition’, and ‘*Blind* condition’) where vocal coordination was not needed (Figure 1d), and one test condition (‘*Blind delayed* condition’) which required vocal communication (Figure 1d). In the ‘Cooperative condition’, the birds were in visual contact (Video SV1) and could freely approach their ends of the string. Synchronization was not necessary if both parrots pulled the string immediately after being released. In the ‘*Delayed* condition’, also with visual contact (Video SV2), access to the string was controlled by trap doors, with a five-second delay if a bird approached and ten seconds if it remained still (Figure 1b). This setup required one bird to wait for its partner in order to successfully retrieve the food. The ‘*Blind* condition’ involved visual isolation using a cardboard wall (Video SV3), allowing the birds to hear each other but not see one another. As the birds were both released at once, they could retrieve the food reward simultaneously without purposeful coordination (Figure 1c). Finally, in the ‘*Blind delayed* condition’, similar to the ‘*Delayed* condition’ but visually isolated, the parrots needed to communicate to synchronize their actions without visual feedback (Figure 1d; Video SV4). All data was recorded with two webcams (Logitech Carl Zeiss Tessar HD 1080p; 25 fps) connected to a computer (using Logitech Webcam Software v2.30). The first camera was 10 centimeters away from the set position of the sliding board and at the same height as the parrots to record a frontal view. The second camera was 20 centimeters from the sliding board and 47 centimeters above the parrots to record a top view.

**Figure 1.**
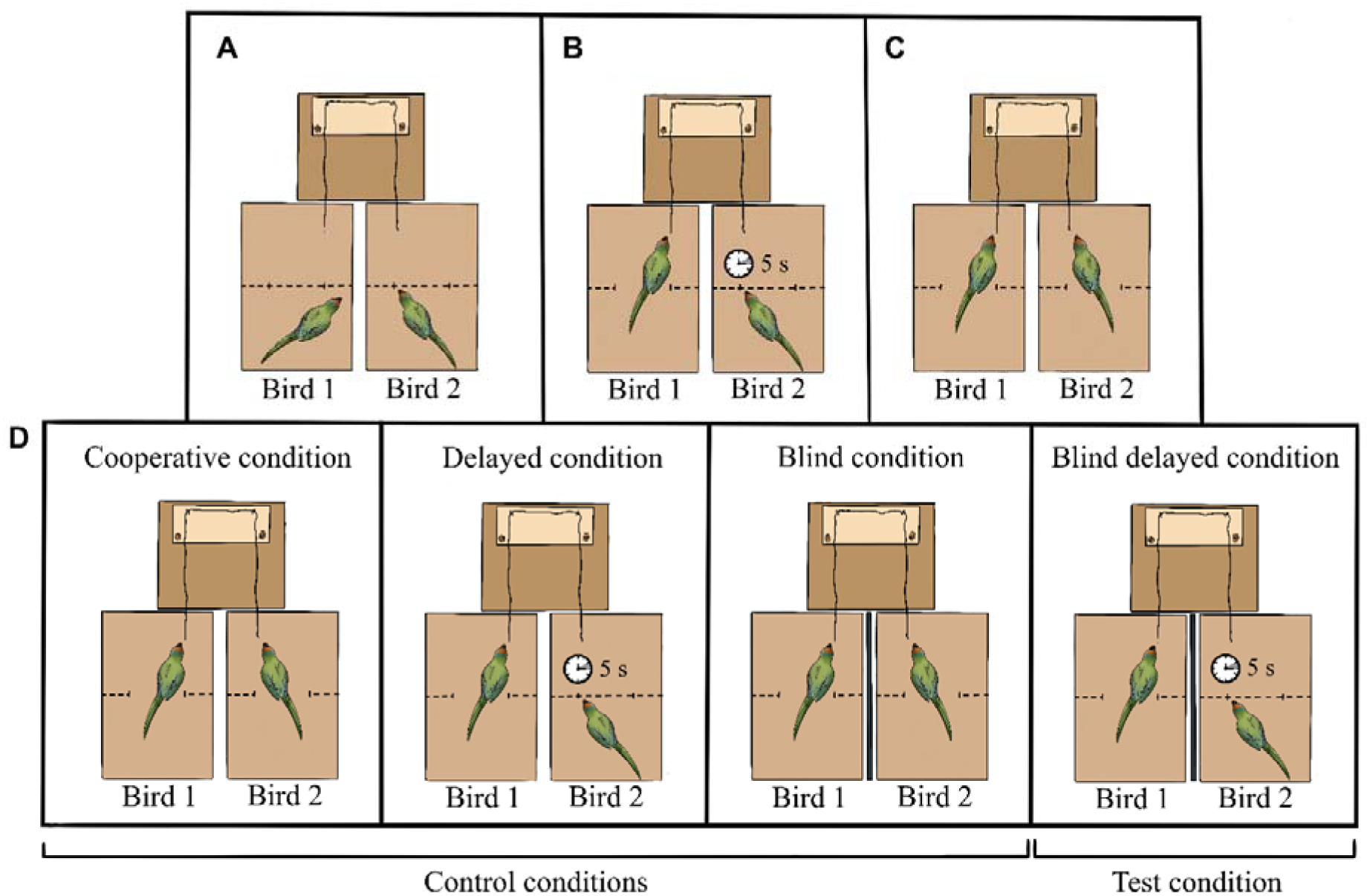
Overview of the control and the test conditions. **(A)** The setup viewed from above with both parrots behind the trap doors. This figure represents the parrots’ position before every trial started. (**B)** This panel shows the position of the birds in the two delayed conditions (‘*Delayed* condition’ and ‘*Blind delayed* condition’). Bird 1 gets access to the string first and needs to wait 5 seconds for the arrival of Bird 2 to solve the task successfully. (**C)** Situation required for successfully solving the task in all conditions: the birds can simultaneously grab the ends of the string and pull the board with the food reward within reach. (**D)** The four experimental conditions, three controls and one test. The thick black line between cages in ‘*Blind*’ and ‘*Blind delayed*’ indicates the visual barrier.

### Setup for recording vocalizations

Vocalizations emitted by the birds during each trial were recorded with a setup consisting of a ½” free-field microphone (Brüel & Kjær, type 4133 with preamplifier type 2669) placed at 15 centimeters above the parrots’ cages and connected to a microphone conditioner (Brüel & Kjær, type 5935; amplifying 20 dB and without filter (Lin)). The frequency response of the entire recording system was 5-20,000 Hz, with a tolerance of ± 0.5 dB. To synchronize video with sounds recorded in this setup, we used a custom-built manual synchronizer that simultaneously emitted a click sound and a red LED light, when activated by the experimenter. The microphone signals were digitized as 16 bit / 44.1 kHz wav files, using *Batsound* version 4.01. Using the videos, we matched each vocalization to the bird producing it. The emitter was clearly identified because they opened their beak to vocalize.

### Testing procedure

Details on the testing procedure can be found in Ortiz et al. (2020) and are summarized in Figure 1. Each bird participated in 20 trials across all pairwise combinations, resulting in a total of 60 trials for conditions without time delays and 120 trials for delayed conditions. In the delayed conditions the birds had different roles: Bird 1 was the one whose door opened first and therefore would have to wait for the second individual, Bird 2, to succeed. Each bird completed 20 trials for each role (Bird 1 and Bird 2) with three partners, explaining the variation in trial numbers.

### Identity and call type clustering

To classify vocalizations, we used hierarchical cluster analysis with the function *hclust* from the *stats* package in R (Müllner, 2013). We used the peak frequency, peak pressure and first quartile for the clustering. All parameters were scaled and transformed. For the transformation we used the Box-Cox power transformation with optimal lambda retrieved from the function *boxcox* from the *MASS* package (Ripley et al., 2013). For each bird and each experiment, the vocalizations were assigned to ten clusters using this technique. For each cluster all the corresponding vocalizations were assigned to nine vocalization types by one of the experimenters, allowing for multiple clusters to be assigned to the same vocalisation type. F1 produced an extra call only in the role as Bird 1, when she had access to the string and was waiting for the arrival of Bird 2. This extra call was named “*readiness-call*” and was separated by the context in which it was emitted.

To test for acoustic differences between the four individuals and between call types, all audio files and call start/end points were imported into *Koe* bioacoustics software (Fukuzawa et al., 2020); koe.io.ac.nz), comprising 2553 calls from 661 trials. (The experiment included a total of 1440 trials, but only 661 trials had calls). In *Koe*, for all calls, the entire suite of available acoustic features was extracted, including all available feature aggregation methods (combining to 7958 features; https://github.com/fzyukio/koe/wiki#extract-unit-features).

To visualize and test whether the four individuals (M1, M2, F1, F2) were acoustically distinct, a linear discriminant analysis (LDA) was performed using the following feature selection:

1. A multinomial generalized linear model (Friedman et al., 2010) via penalized maximum likelihood was fit to all acoustic features (i.e., both those extracted using *Koe* and those from R) to find the most relevant acoustic measures (81 features were selected; Table S5) using the R package *glmnet* (Hastie & Qian, 2014). The regularization path for the LASSO penalty was chosen using 10-fold cross-validation.
2. The features selected by the LASSO were ranked using correlation-adjusted t-score (Ahdesmäki & Strimmer, 2010) and the final number of features was chosen by maximizing leave-one-out cross-validation success.

The features from (2) were then used in an LDA to find the plane that best discriminated between the four individuals (see Figure 5B). Leave-one-out cross-validation correctly assigned calls to the four individuals in 83% of cases.

To visualize and test whether F1’s different vocalization types were acoustically distinct (Figure 5C), the same LDA procedure was used to find the plane that best discriminated between call types for the F1 individual (see Figure 5C). Based on 140 selected acoustic features (Table S6), leave-one-out cross-validation, correctly classified 87% of calls across the 9 call types produced by F1, and correctly identified the *readiness call* in 84% (32/38) of cases.

### Data analysis

Data were analysed using generalized linear mixed models (GLMMs) and generalized linear models (GLMs) in R (R Core Team, 2013). GLMMs were fitted using the *lme4* package (Bates et al., 2015), and parrot pairs performing the trials were included as random.

#### Calling during the four experimental conditions

To test for differences in call numbers between control and test condition, we tested if the probability that a bird called during a trial differed between the four experimental conditions using a generalized linear mixed model:

Call∼Condition+(1∣Pair)

We tested the difference in the number of quiet trials by analysing if the parrots’ probability of vocalizing differed between conditions. We used a generalized linear model with a binomial distribution (Table S4). We included condition, trial, and pair as fixed effects in the model:

Call∼Condition+Trial+Pair

The duration of critical part of the trial where Bird 1 is released and waiting for Bird 2 was so short (∼ 5 s) that the number of vocalizations, which the bird could emit was limited for all conditions except ‘*Cooperative* condition’. We therefore decided to analyze vocal activity during a trial as presence or absence of vocalizations. Despite the short durations, the statistical analysis accounted for the variation in duration by including the logarithm of the duration of the periods as offsets. However, as conditions differed significantly, we decided to analyse the other parameters (ie. trial and pair) within each experiment. For these models we also used a generalized linear model with a binomial distribution.

We analysed if the different roles (Bird 1 and Bird 2; Figure 1) affected the vocal activity (measured as number of calls) during the ‘*Delayed*’ and the ‘*Blind delayed*’ conditions. Here we used a generalized linear mixed model with a Poisson distribution to account for the repeated measures structure since both birds in their different roles contributed with vocalizations. The model included role (i.e., Bird 1 or Bird 2), pair, and success. We used the trial number to specify the repeated measures structure of the data set. The models assumed a first-order autoregressive covariance matrix as the response might depend on the previous trial and whether they have learned what to do in the task:

calls∼role+pair+success+(1∣trial)

We used least square mean differences for all post hoc comparisons. We back-transformed least square mean estimates using exp(estimate) and back-transformed the standard error by exp(estimate)*stderr. We used *Proc genmod* and *Proc glimmix* in SAS vers. 9.4 to conduct the analyses.

Lastly, to explore if the presence or the absence of calls predicted the outcome of the trial in each experimental condition, we built a generalized model with a binomial family where our response variable was the trial outcome (success or failure) and our explanatory variable was call presence or absence (binary).

- utcome∼call_presence

#### Differential use of call types

To assess the differential use of call types among experimental conditions, we investigated the diversity in call types between the four conditions using the Simpson diversity index (Figure S3). During all four experimental conditions, all parrots’ calls were then analysed and separated by individuals in clusters (Figure 5A & B). To compare the types of calls between experiments, Simpson’s Diversity index was calculated using the package *vegan* (Oksanen et al., 2013). Simpson’s Diversity Index quantifies biodiversity by measuring the diversity of species in a community. In this case it was used for measuring diversity of calls accounting for the number of different call types in each experimental condition and the call type evenness, or how evenly call types are distributed among trials. The index was used to measure the diversity which considered the number of call types, as well as the relative abundance of call types. To examine whether call diversity varied across experimental conditions, we conducted a Kruskal-Wallis test (Kruskal & Wallis, 1952) followed by pairwise Wilcoxon tests with Bonferroni correction. These non-parametric tests were used due to the non-normal distribution of diversity indices. Additionally, we used a generalized linear model (GLM) to assess the effects of phase and experimental outcome on vocalization type. The model was formulated as:

type∼phase+outcome

To account for potential random effects of pair identity, we conducted an analysis of variance (ANOVA) on a linear mixed-effects model (LME) with pair identity as a random factor.

To analyse the use of different call types in relation to experimental conditions, including phase, experiment, and outcome, a multinomial logistic regression was performed. This statistical model was chosen because the response variable, "call type," is categorical with more than two possible outcomes (e.g., aggressive, contact, etc.), necessitating the use of multinomial logistic regression as the appropriate GLM for this type of data. The model was specified as follows:

type∼phase+experiment+outcome

The model was fitted using the *multinom* function from the *nnet* package in R. This approach allowed us to assess the relationships between the categorical call types and the predictors of phase, experiment, and outcome.

Chi-square tests were performed for both overall and per-experiment analysis to analyse the relationships between call types, the different phases during a trial and the outcome of the trial. We conducted post-hoc analysis using adjusted standardized residuals and Fisher’s exact tests to determine which specific combinations of call type-trial phase and call type-trial outcome are significantly different from expected frequencies.

Lastly, a logistic distribution was employed to model the probability of succeeding in the trial when the *readiness-call* type was employed.

#### Convergence of calls

To test if birds converged their vocalizations, we used spectrographic cross-correlation with the function *xcorr* from the *warbleR* package (Araya-Salas & Smith-Vidaurre, 2017). This gives a similarity score between each set of vocalizations. If a bird’s vocalization was more similar to the latest vocalization of the other bird compared to the latest vocalization of itself, we scored that vocalization as converging towards its partner’s vocalization. If each bird vocalized at least twice in a trial, we could determine if the birds were overall converging or diverging their vocalizations. If more than 2/3 of the vocalizations in a trial were converging, we classified that trial as convergent. Otherwise, it was classified as non-convergent.

We tested the probability of occurrence of vocal convergence given the observed outcome (success or failure) in a logistic mixed model, where the experimental outcome was included as a fixed effect and the pairs of parrots as random effects.

## Results

### Calling during the four experimental conditions

The likelihood that the parrots vocalized during trials differed significantly between experimental conditions (Figure 2a & b; Table S2, GLMM: F_3,701_ = 10.0, p < 0.001). During the ‘*Blind delayed* condition’ there were significantly more trials with vocalizations, compared to the other three control conditions where we found more quiet trials (Figure 2b; Table S3. Least square means differences (LS means: t_701_ ≥2.35, p < 0.019)). The likelihood of vocalizing during trials did not differ significantly for the control conditions, although the ‘*Cooperative* condition’ tended to show higher likelihood of vocalization than both the ‘*Blind*’ and the ‘*Delayed* condition’ (Table Q2, Least square mean differences, Blind: t_701_=1.81, p=0.07; Delayed: t_701_=1.69, p=0.092). The ‘*Blind*’ and the ‘*Delayed*’ conditions did not differ significantly (Table Q2, least square means differences t_701_=0.48, p=0.632).

**Figure 2.**
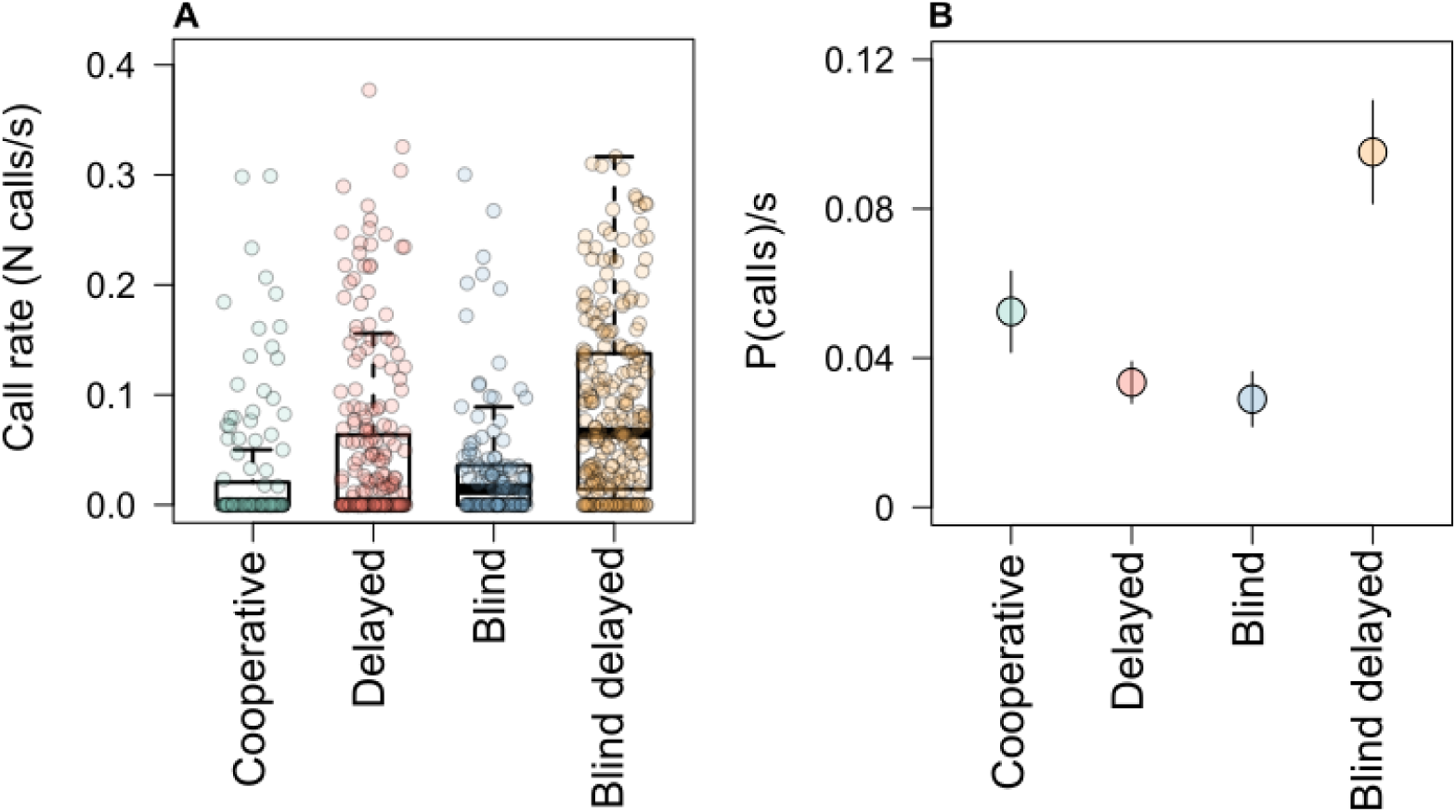
Calling during the four experimental conditions. **(A)** Boxplot of the call rate (number of calls per second) during the four experimental conditions. **(B)** Plot of the GLMM model showing the probability of calls per second during the four experimental conditions. The results of the model are shown in the four differently coloured dots, and the lines show the standard deviation.

**Figure 3.**
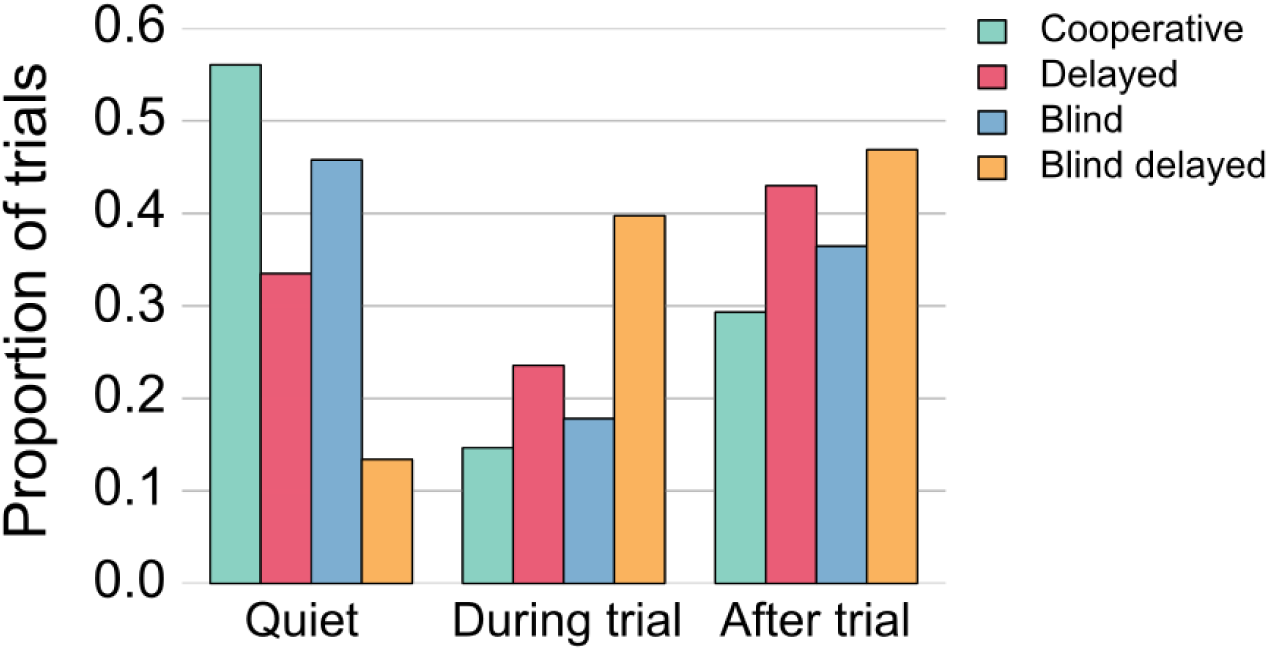
Vocalizations during each trial type in each experiment. Bar plot of the proportion of quiet trials (no calls emitted during the entire trial) in the first set of columns. The proportion of trials with vocalizations only during the part of the trial where the birds need to coordinate (Figure 1B & C) are illustrated in the middle set of columns and the proportion of trials where there were only vocalizations once the trial ended are on the last set of columns. Each of the four experimental conditions are illustrated by the different colours.

We also found differences in calling vs. no-calling related to the success or failure of the trials. When the birds were visually isolated and released at different times in the ‘*Blind delayed* condition’, calling (as opposed to non-calling) significantly increased the number of successful trials (GLM: df = 239, z-value = 3.75, p < 0.001), whereas calling did not correlate with trial success for the other conditions (‘*Delayed* condition’ GLM: df = 239, z-value = -0.82, p = 0.41; ‘*Blind* condition’ GLM: df = 117, z-value = 1.22, p = 0.22).

During the ‘*Blind delayed* condition’ most of the calls were emitted from the moment when Bird 1 arrived at the string and until the door for Bird 2 opened (Figure 4). The other three experimental conditions did not differ significantly, although the ‘*Cooperative* condition’ tended to show higher likelihood of vocalization than both the ‘*Blind* condition’ and the ‘*Delayed* condition’ (Figure 2b; Table S3, LS means, Blind: t_701_ = 1.81, p = 0.070; Delayed: t_701_ = 1.69, p = 0.092), whereas the likelihood of vocalizations during the ‘*Blind*’ and the ‘*Delayed*’ conditions did not differ significantly (Figure 2b; Table S3, LS means: t_701_ = 0.48, p = 0.632).

**Figure 4.**
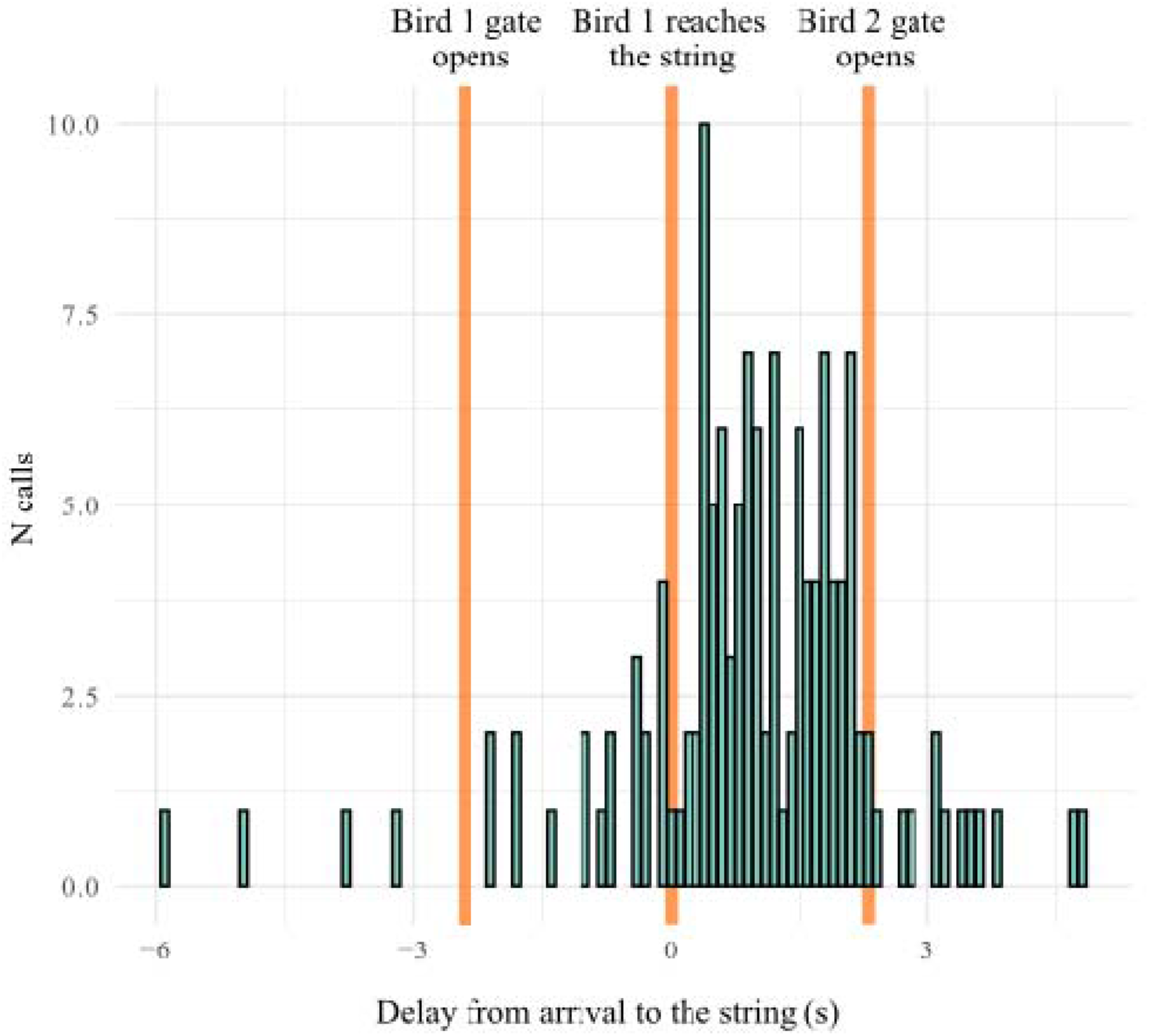
Call timing across all trials in the ‘*Delayed’ and the ‘Blind delayed’* conditions. Time between all calls produced at the different stages of a trial where coordination was needed (marked with orange vertical lines). Zero represents the moment that Bird 1 arrive at the string. The two other markings on the x-axis are when the door for bird 1 opens (at -2.5 seconds) and when the door for bird 2 opens (at +2.5 seconds). On the y-axis we have the raw number of calls emitted in each 0.1 second time interval.

### Differential use of call types

Differences in the four individuals’ calls were found when analysing their entire repertoire (Figure 5A & B). The call types of the two females were largely distinct in acoustic space, whereas the two males could not be easily distinguished from each other (Figure 5B, Table 1). We identified a call type produced only by F1 (‘readiness-call’; Video SV5). She started presenting this call when she was the first to arrive at the string (i.e., in the role as Bird 1) and had to wait for the arrival of her partner (Bird 2). This call was always emitted right after her arrival at the string and was never produced in any of the other three conditions. The readiness-call was acoustically distinct, correctly discriminated from F1’s other call types 84% of the time, based on LDA ’leave-one-out’ cross-validation (Figure 5B & C). When comparing successful trials with failed trials of F1 as Bird 1 in a logistic mixed effects regression model, F1 was 84% (CI 66.4-93.6%) more successful in trials where she had emitted the readiness-call than in those where this call was absent.

**Figure 5.**
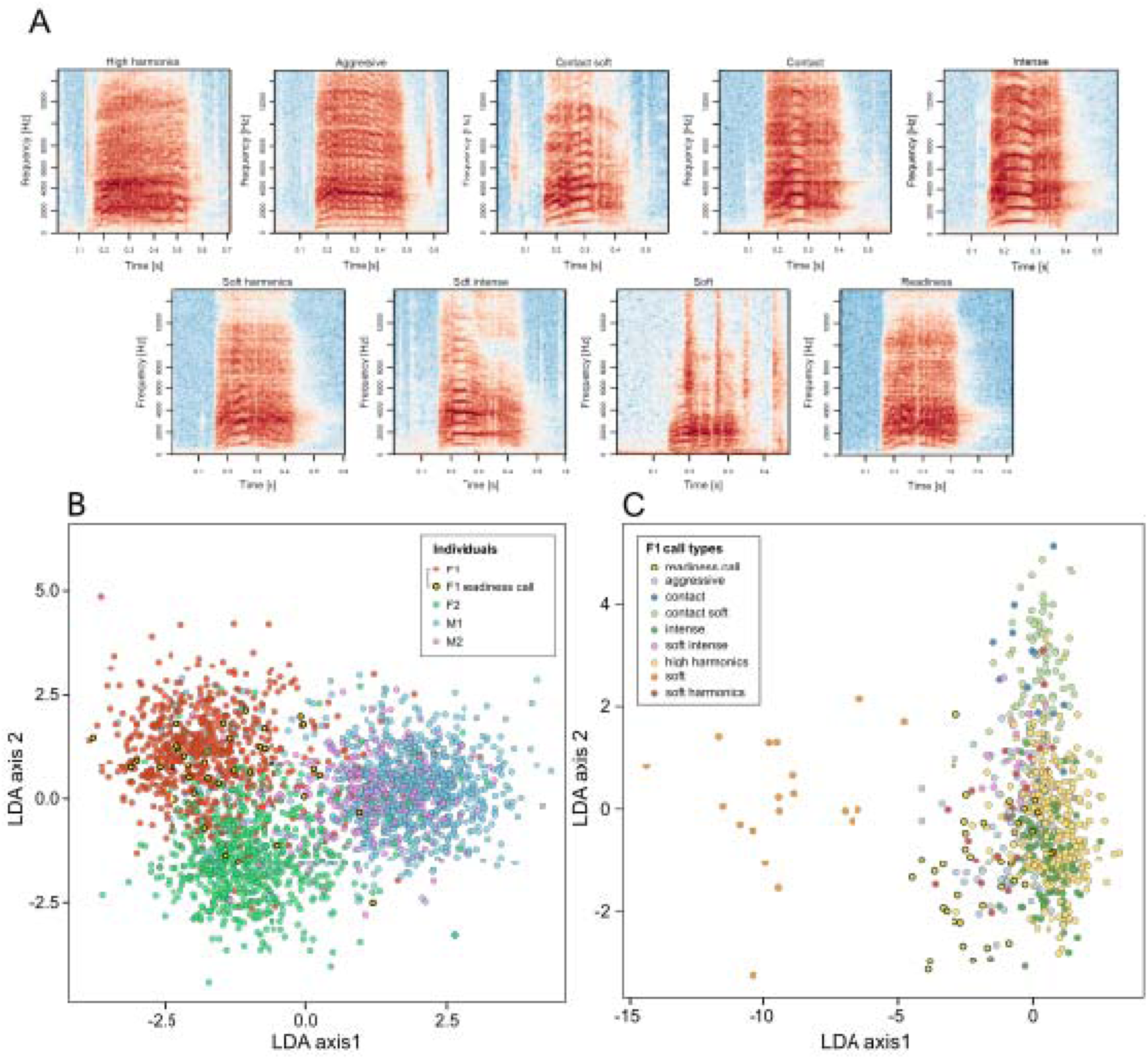
Details of all call types. **(A)** Spectrogram for the nine call types shared by all four birds. **(B)** Two-dimensional linear discriminant analysis (LDA) of calls produced by the four parrot individuals. Each colour represents a different parrot (M1, M2, F1 and F2). The LDA maximizes separation of the four individuals based on acoustic measurements, as detailed in Methods (under ‘Identity and call type clustering’; see Table S5 for list of measurements). The F1 ‘readiness call’ type is highlighted with yellow points. **(C)** Differential use of call types. Two-dimensional linear discriminant analysis (LDA) of all calls produced by female F1. The LDA maximizes separation of F1’s nine call-type clusters based on acoustic measurements, as detailed in Methods (under ‘Identity and call type clustering’; see Table S6 for list of measurements). LDA leave-one-out cross-validation correctly classified 87% of calls across the 9 call types produced by F1, and correctly identified the readiness call in 84% (32/38) of cases.

**Table 1.** Confusion matrix of calls by the four individuals (F1, F2, M1, M2) based on LDA leave-one-out cross-validation. The LDA correctly classified individuals in 83% of runs, based on acoustic properties (See Methods under ‘Identity and call type clustering’).

|  |  | Predicted |  |  |  |
| --- | --- | --- | --- | --- | --- |
|  |  | F1 | F2 | M1 | M2 |
| Actual | F1 | 602 | 71 | 29 | 10 |
|  | F2 | 61 | 582 | 41 | 9 |
|  | M1 | 35 | 22 | 747 | 40 |
|  | M2 | 9 | 20 | 78 | 197 |

For ‘*Cooperative* condition’, five call types were identified for all birds. For ‘*Delayed* condition’ and ‘*Blind* condition’ we found eight call types, and for the last condition, ‘*Blind delayed*’, we identified nine call types, of which eight calls were shared between all individuals (Figure 5C & Figure 6A). The Kruskal-Wallis rank sum test revealed a significant difference in alpha diversity across experimental groups (χ² = 14.946, df = 3, p = 0.0019). Post hoc pairwise Wilcoxon rank sum tests with Bonferroni correction indicated that there was a higher diversity in calls during ‘*Blind delayed*’, where acoustic communication could help to coordinate the string-pulling behaviour (pairwise non-parametric Wilcoxon test: experiment 3-4, df = 3, p = 0.002, all other pairwise comparisons p > 0.1; Figure S3). A higher diversity of calls could be a result of a more complex communication when the birds require additional cues or information in order to successfully coordinate their behaviour to achieve the common goal of bringing the food reward into reach.

**Figure 6.**
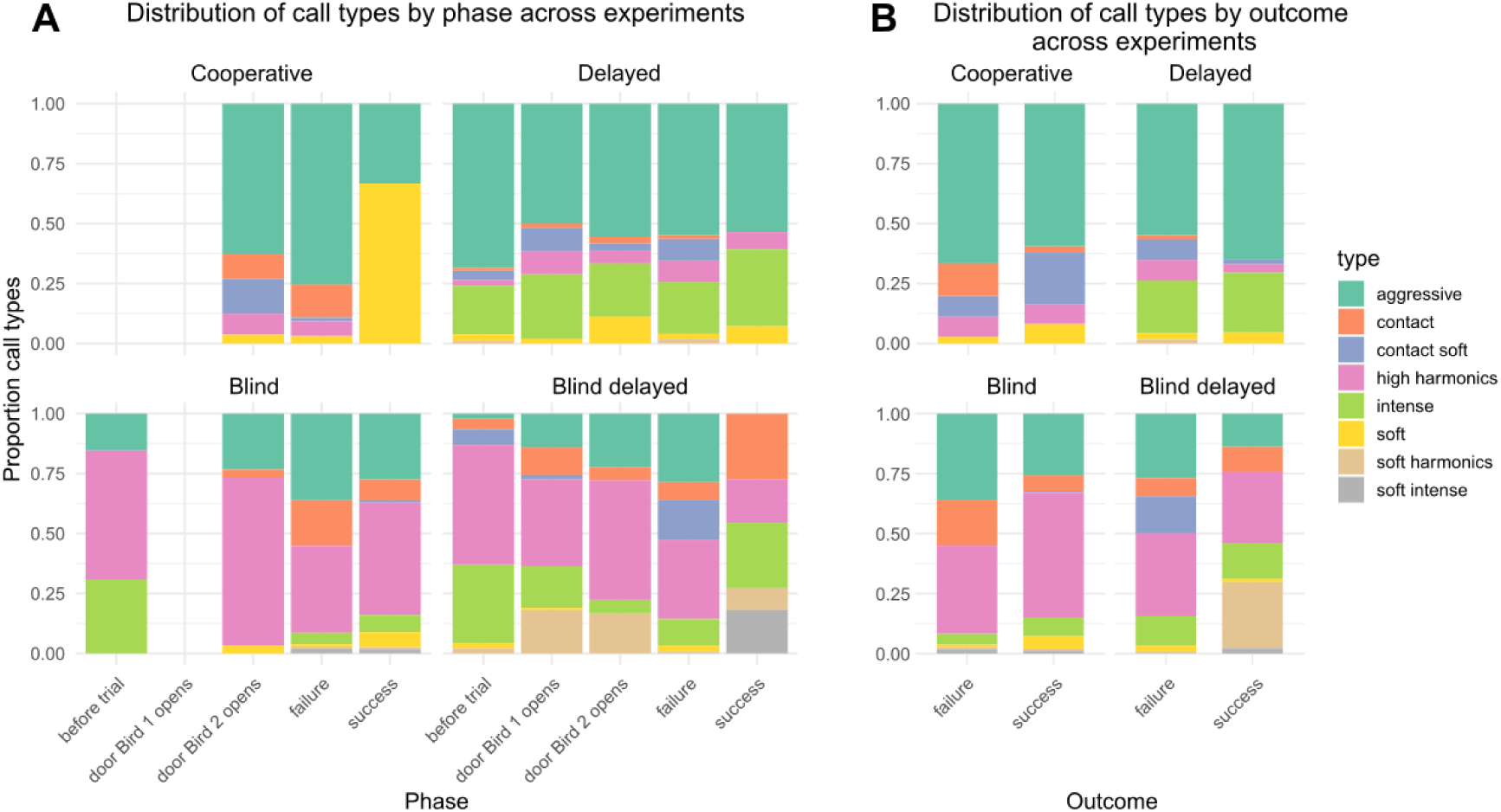
Distribution of call types across experimental phases and outcomes in the different experimental conditions. The y-axis represents the proportion of each call type, with different colors indicating different call types as shown in the legend. The stacked bar format allows for visualization of the relative frequency of each call type within each phase or outcome category. **(A)** Proportional distribution of different call types (aggressive, intense, soft, soft intense) across different experimental phases (before trial, door Bird 1 opens, door Bird 2 opens, failure, success). Data is provided by each experimental condition. **(B)** Proportional distribution of the same call types across trial outcomes (failure, success). Data is provided by each experimental condition.

When looking at all conditions together, both the Chi-square tests and the multinomial logistic regression showed significant relationship between the phase of the trial and the call type (χ²=296.81, df = 28, p-value = 3.56e-51; Figure 7A). We also found a significant relationship between the call type and the outcome: successful or failed trials (χ²=86.023, df = 7, p-value = 8.092e-16; Figure 7B). In failure outcomes, contact soft calls are used significantly more often than in successful outcomes (χ² = 17.80454, df = 1, p-value = 2.448e-05; Figure 7B).

**Figure 7.**
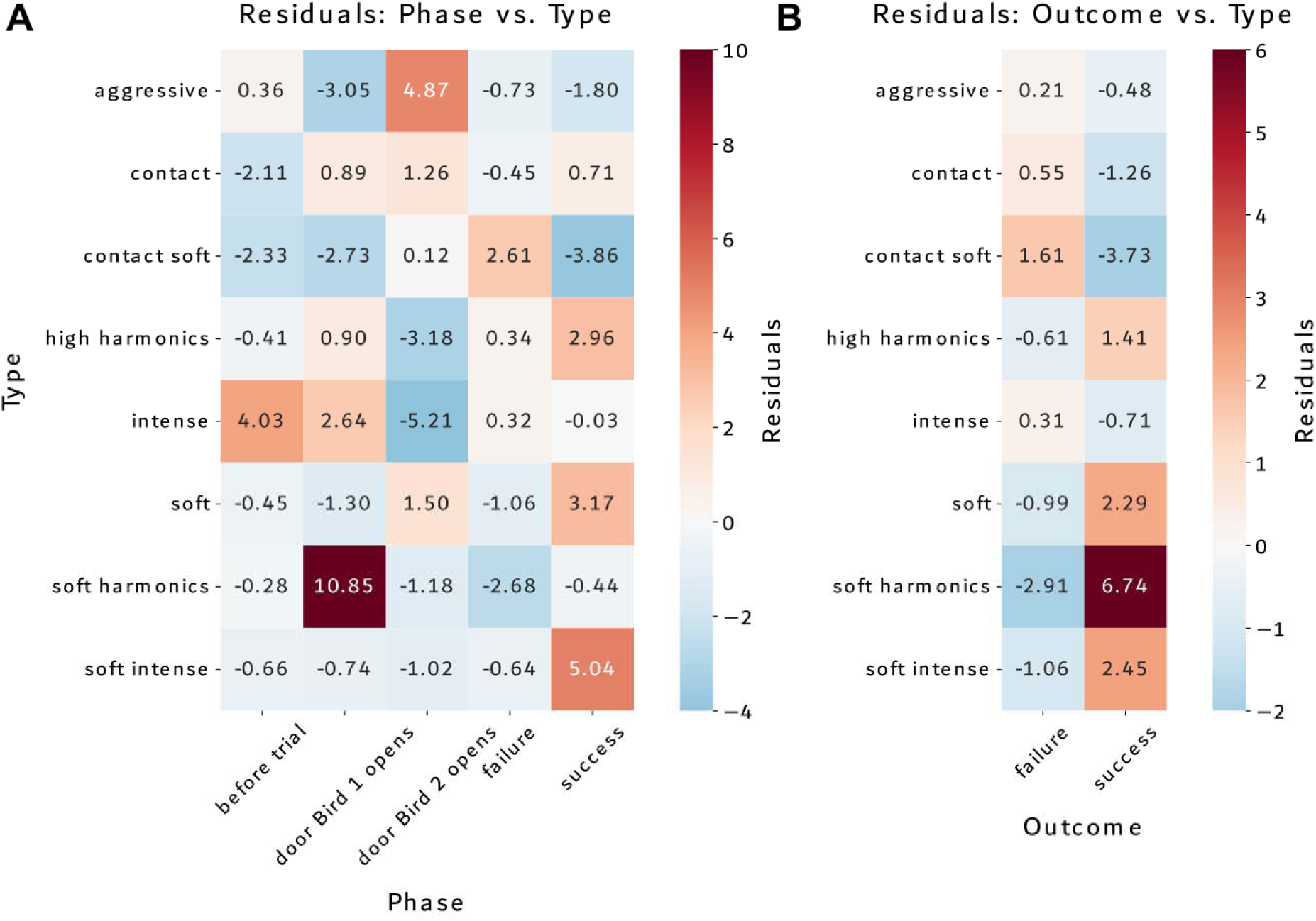
Standardized residuals from chi-square analysis to compare call types. The heatmaps display the standardized residuals, where positive values (red) indicate higher observed frequencies than expected by chance, and negative values (blue) indicate lower frequencies than expected. Colour intensity corresponds to the magnitude of deviation from expected frequencies. Cell values show the exact standardized residual values. Horizontal axis labels indicate the different phases or outcomes, while vertical axis labels show the behavioural types. **(A)** Residuals to compare call types in each trial phase considering all experimental conditions. **(B)** Residuals comparing call type and trial outcome across all experimental conditions.

Looking at each experimental condition separately, we found different patterns in how the parrots used call types. In the ’*Cooperative* condition’, parrots’ clearly changed which calls they used depending on what part of the trial they were in (χ² = 39.999, df = 8, p = 3.205e-06) and whether they succeeded or failed (χ² = 18.501, df = 4, p = 0.001). In the ’*Delayed* condition’, they didn’t really change their call types during different parts of the trial (χ² = 28.994, df = 24, p = 0.22), but they did use somewhat different calls for successful versus unsuccessful outcomes (χ² = 14.329, df = 6, p = 0.026), though this difference wasn’t as strong as in other conditions. When animals couldn’t see each other in the ’*Blind* condition’, they showed some differences in which calls they used during different trial parts (χ² = 39.535, df = 21, p = 0.008) and for different outcomes (χ² = 19.455, df = 7, p = 0.007). The robust differences appeared in the ’*Blind delayed*’ condition, where animals strongly varied their call types both across different trial parts (χ² = 344.75, df = 28, p < 2.2e-16) and depending on whether they succeeded or failed (χ² = 252.43, df = 7, p < 2.2e-16).

We also found that there were significant differences in the distribution of call types in all conditions for the phase when they would need to coordinate, when Bird 1 is released, the waiting time, when Bird 2 is released and until the trial failed or succeeded (Chi-square statistic: 286.754, p-value: 2.53e-50; Figure 1B & C). Looking at each call type specifically during this trial phase, there were significantly more *Aggressive* calls during ‘*Cooperative* condition’ than ‘*Blind delayed* condition’ (χ² = 79.37, df = 1, p-value = 5.15e-19) while there were significantly more *Soft harmonics* and *High harmonics* during the ‘*Blind delayed* condition’ compared to the other three control conditions (χ² = 44.93, df = 1, p-value = 2.04e-11; χ² = 47.23, df = 1, p-value = 6.31e-12). When we analysed the call type, during the part of the trial where birds need to coordinate and the outcome (failure vs. success), we found a significant relationship between call types and trial outcome for the ‘*Blind delayed* condition’ (χ² = 33.22, df = 6, p-value = 0.09e-5; Figure 8).

**Figure 8.**
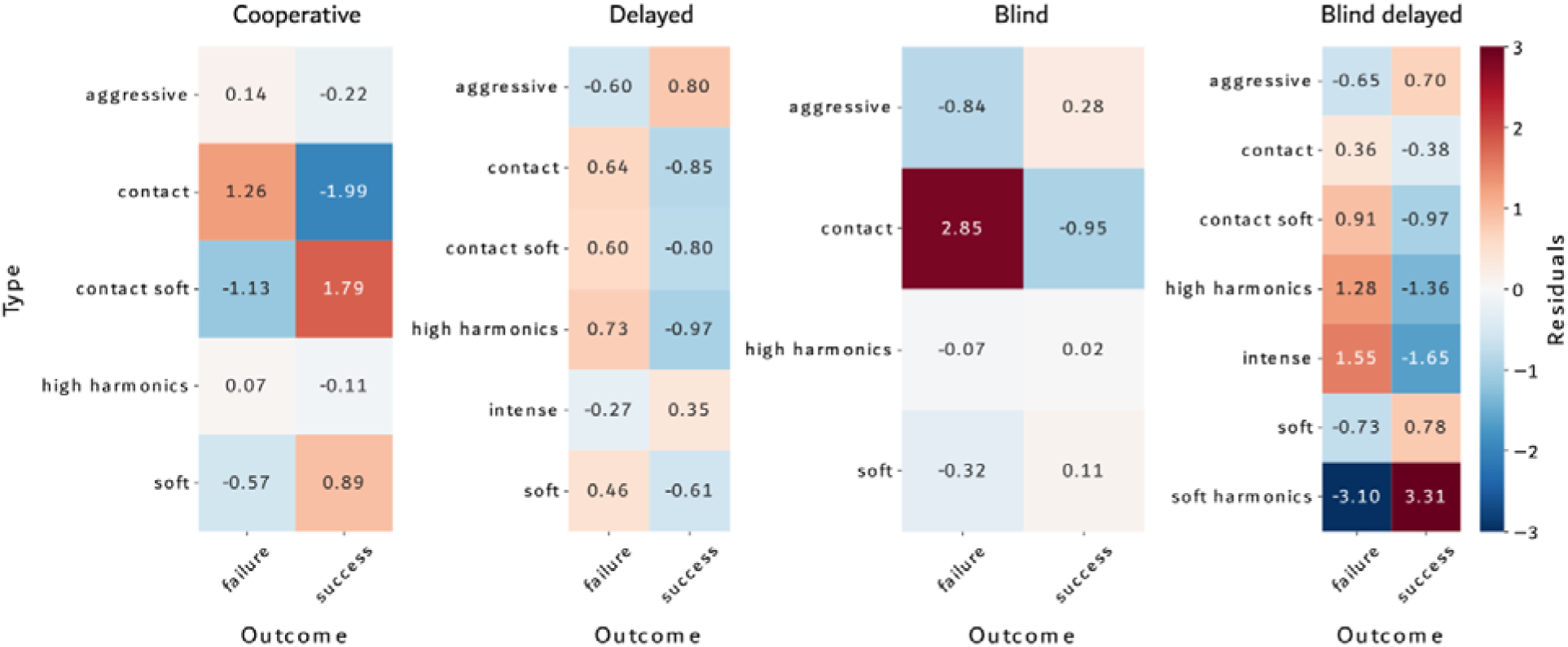
Comparison of standardized residuals across the four experimental conditions (cooperative, delayed, blind and blind delayed). The graph shows the relationship between trial outcome and call type only in the experimental phase where coordination is required. Each panel represents a different experimental condition, with standardized residuals displayed in a heatmap format (scale: -3 to 3). Red colours indicate higher observed frequencies than expected by chance, while blue colours indicate lower frequencies. Cell values represent standardized residuals from chi-square tests, with more extreme values indicating stronger deviations from expected frequencies. The colourbar on the rightmost panel provides a reference scale for all plots. This visualization allows direct comparison of type-outcome associations across experimental conditions.

The Fisher’s exact test results for individual call types in the ‘*Blind delayed* condition’ showed that *Soft harmonics* had a significant positive association with success (OR = 19.71, p-value = 0.017e10−6; Figure 8). *High harmonics* and *Intense* calls showed a significant negative association with success (OR = 0.43, p-value = 0.023; OR = 0.28, p-value = 0.019; Figure 8). *Aggressive* and *Contact* calls, however, showed no significant association with success (OR = 1.64, p-value = 0.348; OR = 0.73, p-value = 0.785; Figure 8) and *Soft* and *Contact soft* calls had too few observations for reliable statistical inference. For the three control conditions, overall Chi-square test for call types vs outcomes in ‘*Cooperative* condition’ showed there are some call types associated with the outcome of the trial (χ² = 11.22, df = 4, p-value = 0.024). Specifically, *Contact soft* calls where significantly associated with both success and failure (OR= 2.30; p-value = 0.028; Figure 8) and *Contact* call was associated with failure (OR= 0.19; p-value = 0.011; Figure 8). The ‘*Delayed* condition’ did not show an association between call type and trial outcome (χ² = 5.40, df = 5, p-value = 0.368; Figure 8). Lastly, the ‘*Blind* condition’ did show an association between trial outcome and call type when taking all call types in the analysis (χ² = 9.89, df = 3, p-value = 0.019; Figure 8) but when performing the Fisher’s exact test for individual call types association, we did not find a specific call type significantly associated with success or failure since individual call types either lacked sufficient variability or sample size was too small to independently show significance, as seen in both Fisher’s exact test and chi-square tests for individual call types.

### Convergence after failure

Taking all conditions together, the peach-fronted conures tended to use rapid convergence more often after failed trials, i.e., when the food reward was not retrieved (GLMM: df = 66, z-value = -1.64, p = 0.099), compared to successful trials.

## Discussion

Our study is the first to examine vocal communication during a cooperative task in a non-mammalian species, where the identity of all the emitters is known. Although our sample size was small, our results strongly suggest that peach-fronted conures can use vocal communication to coordinate individual behaviour in a cooperative task – the first evidence for vocally coordinated task-solving in a non-mammal. Vocal communication within dyads occurred mostly when visual feedback was not available, and the birds had to coordinate since they were not released simultaneously. The conures were able to coordinate their behaviour to solve the task in all four experimental conditions. The previous behavioural analysis (Ortiz et al., 2020) showed that, in many of the delayed trials, the individual in the role of Bird 1 did not use non-vocal cues to anticipate the arrival of Bird 2, such as the sound of the footsteps from the other bird, or the string moving. Rather, the differences found both in the call rate, the number of quiet trials between ‘*Blind delayed*’ and the three control conditions where significantly more quiet trials were found in the control conditions when coordination was not required to solve the task. This, together with the likelihood of succeeding when calls were present only in this condition, are strong indicators that the parrots used vocalizations to synchronize their behaviour to solve the task. Our results revealed significant differences in the distribution of call types between the *Blind Delayed* condition and the three control conditions, both overall and specifically at the critical moment when both birds needed to coordinate. Notably, the call type *Soft Harmonics* was the only call observed during the *Blind Delayed* condition, occurring precisely at the key coordination moment of each trial. Moreover, the presence of this call type was a predictor of success in the *Blind Delayed* condition, providing evidence of non-random relationships between call types, trial phases, and trial outcomes.

The strength of these relationships varied across experimental conditions. Only in the *Blind Delayed* condition (where parrots relied on vocalizations to coordinate) did we observe a clear association between a single call type (*Soft Harmonics*) and trial success. While control conditions exhibited some patterns, such as *Contact Calls* being associated with both success and failure, these associations were not as distinct as in the test condition. Given the pronounced patterns observed in the *Blind Delayed* condition, our findings suggest that this experimental setup influenced the parrots’ communication strategies.

In a previous study (Tassin de Montaigu et al., 2020), blue throated macaws were not able to coordinate their behaviour to solve the ‘*Delayed* condition’ with visual feedback, and their vocal activity did not vary between conditions. Our parrots seemingly understood the need for a partner in the ‘*Delayed* condition’, which could be the reason for their use of vocal communication. In the wild, conures are highly social and appear to use vocalizations to negotiate and share information about food sources when flocks meet (Bradbury & Balsby, 2016). That our experimental conures used vocal communication to solve the cooperative task is plausible given the life history of this species. Individual differences regarding contact calls have been reported previously by Thomsen et al., (2013) but in the present experiment, we detected individual differences also when considering the whole repertoire of call types. We also found a call type produced only by F1 (‘readiness-call’). She emitted this call always right after her arrival to the string and only in the test condition (‘*Blind delayed*’). F1 was also the most successful parrot in solving the ‘*Blind delayed* condition’, where vocal communication was needed.

In our study, we often observed rapid call convergence after trials had ended. Rapid vocal convergence occurs when an individual produces vocalization with acoustic features that are increasingly similar to calls of another individual that is present. Previous work has shown rapid call convergence in the wild for the closely related to our test model, orange-fronted conures (*Eupsittula canicularis*). Balsby and Scarl (2008) suggested that rapid vocal imitation may serve to mediate transient affiliation with an individual or a group, to negotiate dominance, or to make group decisions. For budgerigars, it has been shown that rapid vocal convergence occurs in males for pair bonding with females, while females did not imitate their mates (Hile et al., 2000). This leads us to suggest that rapid call convergence may also be used to reconcile the partners after unfavourable or frustrating experiences (Videos SV6 and SV7 for convergence example; See Video SV8 for non-convergence example).

Non-vocal communication has been suggested as a means of solving similar tasks, for instance glancing rates (Chalmeau & Gallo, 1996; Hattori et al., 2005; Mendres & de Waal, 2000) and gestural pointing (Melis & Tomasello, 2019). However, only two previous studies had no visual feedback (King et al., 2021; Mendres & de Waal, 2000): the performance of chimpanzees during the visually isolated condition dropped considerably but their vocal communication was not analysed (Mendres & de Waal, 2000). For dolphins (King et al., 2021), possible differences in communication and performance between trials with and without visual feedback were not analysed.

Parrots demonstrate a range of sophisticated vocal behaviors that enable complex social interactions, particularly through vocal learning and imitation (Pepperberg, 1990, 2002). Unlike primates, which use a combination of gestures and vocalizations, parrots rely predominantly on vocal communication due to their limited gestural repertoire (Moura et al., 2014). Most research on parrot vocalizations has been based on observational and playback studies (Balsby & Bradbury, 2009; Balsby & Scarl, 2008; Bradbury et al., 2001; Wright et al., 2003), yet their communication system appears to be highly structured and flexible.

For example, peach-fronted conures and related species use individually distinctive contact calls, which are believed to play a role in social coordination (Thomsen et al., 2013). Field studies on orange-fronted conures have shown that these parrots can quickly imitate the calls of specific individuals (Balsby et al., 2012; Vehrencamp et al., 2003), suggesting that they use vocal mimicry to establish direct communication within their social networks—similar to the way bottlenose dolphins (*Tursiops* sp.) address one another by copying signature whistles (King & Janik, 2013). This ability to imitate vocalizations may serve various functions, including selecting social partners, coordinating group movements, negotiating leadership, or signaling food locations (Balsby et al., 2012; Bradbury & Balsby, 2016). These findings indicate that parrot vocal communication is not only adaptable but also plays a key role in shaping their social interactions.

Our results strongly suggest that the birds used vocal communication to coordinate and solve the cooperative task. First, the number of calls was significantly higher when they were visually isolated. In contrast, when visual contact was available the number of quiet trials was also higher (Figure 3). In particular, we observed one specific call, referred in our study as *Soft harmonics* strongly associated with the critical phase of the trial where parrots need to coordinate only in the experimental condition ‘*Blind delayed*’, and the occurrence of this call was also associated with a successful outcome of the trial. Our results highlight the birds’ use of vocalizations to adapt their behaviour to different task demands. The birds’ success rate was expected to drop once the individuals were visually isolated but, in our experiment, it did not drop between the ‘*Delayed* condition’ and the ‘*Blind delayed* condition’ (Ortiz et al., 2020), in contrast to the results of experiments with capuchin monkeys (Mendres & de Waal, 2000), and for individual F1 the success rate even increased.

The discovery of vocally mediated coordination in conures raises the exciting possibility that this trait is widespread and warrants behavioural experiments regarding vocal coordination in many more taxa.

## Acknowledgements

We thank Simeon Q. Smeele for help with writing the code to extract and analyse all calls from the recordings, and his help on writing and improving the manuscript. Arturo Torres Ortiz for helping with analysis and graph design. We also thank Kasper Fjordside for helping with the setup electronics and Angelica Munteanu and Alejandro Corregidor for assisting in parts of the data collection. We are grateful to Magnus Wahlberg, Coen Elemans and Anastasia Krasheninnikova for comments to improve the manuscript. This project was funded by the Danish Council for Independent Research L Natural Sciences through grants to Ole Næsbye Larsen [DFF – 1323-00105].

## Author contributions

Conceptualization, S.T.O. and O.N.L.; Methodology, S.T.O., T.J.S.B. and O.N.L.; Investigation, S.T.O., T.J.S.B. and O.N.L.; Formal analysis, S.T.O., W.W., M.P., T.J.S.B. and O.N.L.; Software, W.W. and M.P.; Writing – Original Draft, S.T.O., T.J.S.B. and O.N.L.; Writing – Review & Editing, S.T.O., W.W., M.P., T.J.S.B. and O.N.L.; Funding Acquisition, O.N.L.; Resources, S.T.O., W.W., M.P., T.J.S.B. and O.N.L.; Supervision, T.J.S.B., and O.N.L.

## Declaration of interests

The authors declare no competing interests.

### Ethics

All applicable international, national, and institutional guidelines for the care and use of experimental animals were followed.

### Data accessibility

Data is available in the Zenodo repository (https://zenodo.org/deposit/6647572).

