## Supplemental Information for "Vocally mediated coordination during a cooperative task in parrots"

**SUPPLEMENTARY MATERIAL**

*Call number, call type and call convergence*

All wav-files (one file per trial) were imported in R (R Core Team (2020). R: A language and environment for statistical computing. R Foundation for Statistical Computing, Vienna, Austria. URL https://www.R-project.org/). Session recordings were only analyzed if the maximum absolute peak-pressure of the uncalibrated signal was above 3000. This number was chosen such that it would be below the amplitude of the soft calls, but above most other noise and ensured that empty recordings were not analyzed. To isolate vocalizations, we used the *autodetec* function from the *warbleR* package. This function implements a bandpass filter, an envelope and detects events that are above a certain percentage of the maximum amplitude in the file. We used the following settings: minimal amplitude to trigger detection = 1% of the absolute maximum amplitude in the file; bandpass filter = 0.5-5 kHz; ssmooth = 1,000; minimal duration = 0.1 s. Each selected vocalization was saved in a separate wav-file (see clips in Figure S2) with the original sampling rate. 0.2 s were added to each end of the bounds to make sure that the entire vocalization was included in the file. The start and end time in the original wav-file were also saved in the new filename. We assessed all files by listening to them and removed files that contained no parrot vocalizations. These first steps were necessary to get rid of the noise (human speech, tapping, etc.).

We imported each of the short sound clips in R for further analysis. To determine the exact start and end of the vocalization we again used *autodetec*, which was now based on the maximum absolute pressure of only the vocalization in question. We used the following settings: minimal amplitude to trigger detection = 15% of the absolute maximum amplitude; bandpass filter = 1-50 kHz; ssmooth = 1000; minimal duration = 0.05 s. If multiple vocalizations were detected, the ones in the 0.2 s tails were excluded. If there were still multiple vocalizations detected, e.g., when two birds were vocalizing at the same time, the vocalizations were excluded because the start and end time could not be reliably determined. For all vocalizations, the following parameters were measured:

1) Peak Frequency (Hz): the frequency with maximum amplitude in a spectrum (hanning window with length = 512) of the vocalization; using the functions *spec* and *fpeaks* from the package *Seewave*
2) Duration (s): the time of the start to end from the *autodetec* function
3) Peak pressure (unit): the maximum absolute amplitude in the time domain
4) First frequency quartile (Hz): using the function *specprop* from the package *Seewave.*

**Table S1.** Confusion matrix of calls by individual F1. Leave-one-out cross-validation overall correctly classified call types in in 87% of cases, and the readiness call in 84% (32/38 cases), on the basis of acoustic measurements (detailed in Methods, under “Identity and call type clustering”; See Table S6 for list of measurements).

|  | |  | **Predicted** | | | | | | | | | |
| --- | --- | --- | --- | --- | --- | --- | --- | --- | --- | --- | --- | --- |
| **Actual** | **Call type** | | | **Aggressive** | **Contact** | **Contact soft** | **Intense** | **Soft intense** | **Readiness-call** | **High harmonics** | **Soft** | **Soft harmonics** |
|  | **Aggressive** | | | 61 | 0 | 0 | 0 | 0 | 2 | 10 | 0 | 0 |
|  | **Contact** | | | 0 | 15 | 0 | 0 | 0 | 0 | 0 | 0 | 0 |
|  | **Contact soft** | | | 0 | 0 | 80 | 2 | 0 | 0 | 10 | 0 | 0 |
|  | **Intense** | | | 0 | 0 | 0 | 76 | 0 | 0 | 18 | 0 | 0 |
|  | **Soft intense** | | | 1 | 0 | 2 | 1 | 24 | 0 | 3 | 0 | 0 |
|  | **Readiness-call** | | | 3 | 0 | 0 | 1 | 0 | 32 | 2 | 0 | 0 |
|  | **High harmonics** | | | 9 | 0 | 10 | 10 | 1 | 2 | 292 | 0 | 1 |
|  | **Soft** | | | 0 | 0 | 0 | 0 | 0 | 0 | 1 | 19 | 0 |
|  | **Soft harmonics** | | | 1 | 0 | 0 | 0 | 0 | 2 | 2 | 0 | 19 |

**Table S2.** Generalized linear mixed model on likelihood of vocalizing in all four conditions during and after trials.

|  | During | | | After | | |
| --- | --- | --- | --- | --- | --- | --- |
| Effect | df | F | p | df | F | p |
| Condition | 3, 701 | 10.00 | <.0001 | 3, 700 | 15.86 | <.0001 |
| Trial | 1, 701 | 23.19 | <.0001 | 1, 700 | 28.84 | <.0001 |
| Pair | 5, 701 | 4.54 | 0.0004 | 5, 700 | 2.34 | 0.0403 |

**Table S3.** Least square mean difference for treatment effect during and after trials. df_during_=701, df_after_=700.

|  | | During | | After | |
| --- | --- | --- | --- | --- | --- |
| Condition | Condition | t | p | t | p |
| Blind | Delayed | -0.48 | 0.6316 | -3.28 | 0.0011 |
| Blind | Blind delayed | -4.11 | <.0001 | -5.02 | <.0001 |
| Blind | Cooperative | -1.81 | 0.0704 | 1.01 | 0.3122 |
| Delayed | Blind delayed | -4.78 | <.0001 | -2.2 | 0.0285 |
| Delayed | Cooperative | -1.69 | 0.0915 | 4.34 | <.0001 |
| Blind delayed | Cooperative | 2.35 | 0.0188 | 6.00 | <.0001 |

**Table S4.** Generalized linear models for each treatment during and after trials. Bold indicates significant differences.

|  | | | During the trial | | | After the trial | | |
| --- | --- | --- | --- | --- | --- | --- | --- | --- |
| Condition | Source | DF | ChiSq | ProbChiSq | Parameter estimates | ChiSq | ProbChiSq | Parameter estimates |
| Blind | Trial | 1 | 2 | 0.157 | -0.065 | 2.16 | 0.141 | -0.051 |
| Blind | Pair | 5 | 18.02 | **0.003** |  | 11.74 | **0.038** |  |
| Delayed | Trial | 1 | 21.02 | **<0.001** | -0.144 | 11.98 | **0.001** | -0.110 |
| Delayed | Pair | 5 | 25.59 | **<0.001** |  | 15.19 | **0.01** |  |
| Blind delayed | Trial | 1 | 0.25 | 0.619 | 0.015 | 0.88 | 0.347 | -0.041 |
| Blind delayed | Pair | 5 | 81.63 | **<0.001** |  | 13.34 | **0.02** |  |
| Cooperative | Trial | 1 | 24.37 | **<0.001** | -0.232 | 1.04 | 0.308 | -0.048 |
| Cooperative | Pair | 5 | 15.24 | **0.009** |  | 22.33 | **<0.001** |  |

**Table S5.** List of the 81 acoustic features used in Linear Discriminant Analysis of the four individuals (Figure 4). After strongly correlated features were removed, this suite of largely Mel Frequency Cepstrum Coefficient-related features (not easily interpretable) was found to be most important for separating individuals. The features were extracted with *Koe* bioacoustics software (feature descriptions available at <https://github.com/fzyukio/koe/wiki#extract-unit-features>) and https://zenodo.org/deposit/6647572).

| **Order of importance for LDA** | **Feature name** |
| --- | --- |
| 01 | mfcc_divcon_7_median.24 |
| 02 | sem |
| 03 | spectral_bandwidth_mean |
| 04 | goodness_of_pitch_begin |
| 05 | mfcc_delta_min.7 |
| 06 | mfcc_delta2_divcon_7_median.68 |
| 07 | mfcc_divcon_7_std.49 |
| 08 | mfcc_divcon_3_median.3 |
| 09 | mfcc_divcon_7_mean.3 |
| 10 | mfcc_divcon_7_median.43 |
| 11 | spectral_bandwidth_median |
| 12 | mfcc_divcon_5_median.3 |
| 13 | mfcc_min.6 |
| 14 | mfcc_delta2_max.19 |
| 15 | mfcc_divcon_5_mean.3 |
| 16 | mfcc_divcon_3_median.43 |
| 17 | mfcc_divcon_7_median.6 |
| 18 | mfcc_delta2_median.3 |
| 19 | spectral_contrast_divcon_7_std.20 |
| 20 | mfcc_median.3 |
| 21 | mfcc_max.4 |
| 22 | mfcc_delta2_divcon_7_median.26 |
| 23 | mfcc_delta_divcon_7_median.18 |
| 24 | mfcc_divcon_3_median.23 |
| 25 | mfcc_delta2_divcon_5_median.117 |
| 26 | mfcc_divcon_7_median.123 |
| 27 | spectral_contrast_divcon_7_median.15 |
| 28 | mfcc_delta2_divcon_5_median.41 |
| 29 | mfcc_divcon_7_mean.24 |
| 30 | mfcc_min.4 |
| 31 | mfcc_delta2_divcon_5_median.115 |
| 32 | mfcc_delta2_divcon_7_median.34 |
| 33 | mfcc_delta2_divcon_5_median.30 |
| 34 | mfcc_mean.3 |
| 35 | mfcc_delta2_divcon_5_median.89 |
| 36 | mfcc_delta2_divcon_7_median.18 |
| 37 | spectral_contrast_divcon_7_median.41 |
| 38 | mfcc_divcon_7_median.83 |
| 39 | mfcc_delta2_min.2 |
| 40 | mfcc_divcon_3_mean.23 |
| 41 | mfcc_delta_divcon_3_median.52 |
| 42 | mfc_divcon_7_median.10 |
| 43 | mfcc_divcon_7_mean.23 |
| 44 | mfcc_delta2_divcon_3_median.47 |
| 45 | spectral_flatness_divcon_7_std.0 |
| 46 | mfcc_delta2_divcon_3_median.27 |
| 47 | mfcc_divcon_3_mean.3 |
| 48 | mfcc_delta2_max.24 |
| 49 | spectral_flatness_divcon_7_std.3 |
| 50 | spectral_flatness_divcon_5_std.0 |
| 51 | mfcc_delta2_divcon_7_std.69 |
| 52 | mfcc_delta2_divcon_3_median.90 |
| 53 | mfcc_begin.3 |
| 54 | mfcc_delta2_divcon_7_std.59 |
| 55 | mfcc_delta2_divcon_7_median.183 |
| 56 | spectral_rolloff_divcon_5_median.0 |
| 57 | mfcc_divcon_7_mean.42 |
| 58 | mfcc_delta2_max.5 |
| 59 | mfcc_delta2_divcon_7_std.75 |
| 60 | mfcc_divcon_5_mean.43 |
| 61 | mfcc_delta_divcon_7_median.24 |
| 62 | average_frame_power_divcon_5_std.0 |
| 63 | mfcc_delta2_divcon_7_std.183 |
| 64 | mfcc_delta_max.14 |
| 65 | mfcc_delta2_divcon_7_mean.26 |
| 66 | spectral_centroid_divcon_5_std.0 |
| 67 | mfcc_delta2_divcon_7_median.160 |
| 68 | spectral_rolloff_divcon_7_median.0 |
| 69 | mfc_divcon_7_std.90 |
| 70 | spectral_contrast_divcon_5_std.26 |
| 71 | mfcc_divcon_7_mean.63 |
| 72 | mfcc_delta_divcon_5_median.1 |
| 73 | spectral_flatness_divcon_7_std.1 |
| 74 | spectral_contrast_divcon_7_median.6 |
| 75 | mfc_divcon_3_median.12 |
| 76 | mfcc_divcon_7_mean.83 |
| 77 | mfcc_delta2_divcon_7_median.138 |
| 78 | mfcc_max.2 |
| 79 | mfcc_divcon_5_mean.24 |
| 80 | spectral_contrast_std.6 |
| 81 | mfcc_delta_divcon_7_median.22 |

**Table S6.** List of the 140 acoustic features used in Linear Discriminant Analysis of the 9 call types given by individual F1 (Figure 5). After strongly correlated features were removed, this suite of largely Mel Frequency Cepstrum Coefficient-related features (not easily interpretable) was found to be most important for separating call types. The features were extracted with *Koe* bioacoustics software (feature descriptions available at <https://github.com/fzyukio/koe/wiki#extract-unit-features>) and https://zenodo.org/deposit/6647572).

| **Order of importance for LDA** | **Feature name** |
| --- | --- |
| 1 | peak |
| 2 | peakpres |
| 3 | Q25 |
| 4 | prec |
| 5 | am |
| 6 | spectral_centroid_divcon_5_std.2 |
| 7 | spectral_contrast_median.5 |
| 8 | spectral_contrast_divcon_3_median.17 |
| 9 | spectral_contrast_divcon_5_median.31 |
| 10 | spectral_contrast_divcon_7_median.1 |
| 11 | spectral_contrast_divcon_7_std.45 |
| 12 | spectral_contrast_divcon_7_std.46 |
| 13 | spectral_contrast_min.4 |
| 14 | mfcc_median.13 |
| 15 | mfcc_divcon_3_mean.57 |
| 16 | mfcc_divcon_5_mean.40 |
| 17 | mfcc_divcon_5_mean.77 |
| 18 | mfcc_divcon_5_mean.93 |
| 19 | mfcc_divcon_7_mean.117 |
| 20 | mfcc_divcon_3_median.53 |
| 21 | mfcc_divcon_5_std.21 |
| 22 | mfcc_divcon_5_std.36 |
| 23 | mfcc_divcon_5_std.63 |
| 24 | mfcc_divcon_5_std.73 |
| 25 | mfcc_divcon_5_std.96 |
| 26 | mfcc_divcon_7_median.63 |
| 27 | mfcc_divcon_7_median.77 |
| 28 | mfcc_divcon_7_median.86 |
| 29 | mfcc_divcon_7_std.51 |
| 30 | mfcc_variance.11 |
| 31 | mfcc_begin.6 |
| 32 | frequency_modulation_divcon_5_mean.1 |
| 33 | frequency_modulation_divcon_7_mean.2 |
| 34 | goodness_of_pitch_divcon_3_mean.1 |
| 35 | goodness_of_pitch_divcon_5_mean.2 |
| 36 | goodness_of_pitch_divcon_5_median.2 |
| 37 | frame_entropy_divcon_3_mean.0 |
| 38 | max_frame_power_divcon_5_std.2 |
| 39 | spectral_flux_divcon_7_std.2 |
| 40 | spectral_skewness_divcon_5_median.1 |
| 41 | spectral_kurtosis_divcon_7_std.5 |
| 42 | harmonic_ratio_mean |
| 43 | harmonic_ratio_median |
| 44 | harmonic_ratio_divcon_5_std.1 |
| 45 | fundamental_frequency_divcon_3_median.1 |
| 46 | fundamental_frequency_divcon_5_std.1 |
| 47 | mfc_mean.8 |
| 48 | mfc_mean.17 |
| 49 | mfc_median.8 |
| 50 | mfc_median.21 |
| 51 | mfc_divcon_3_mean.51 |
| 52 | mfc_divcon_3_mean.58 |
| 53 | mfc_divcon_5_mean.27 |
| 54 | mfc_divcon_5_mean.41 |
| 55 | mfc_divcon_5_mean.101 |
| 56 | mfc_divcon_7_mean.55 |
| 57 | mfc_divcon_7_mean.61 |
| 58 | mfc_divcon_7_mean.127 |
| 59 | mfc_divcon_7_mean.141 |
| 60 | mfc_divcon_7_mean.257 |
| 61 | mfc_divcon_3_median.9 |
| 62 | mfc_divcon_3_median.64 |
| 63 | mfc_divcon_3_median.89 |
| 64 | mfc_divcon_3_std.14 |
| 65 | mfc_divcon_3_std.50 |
| 66 | mfc_divcon_3_std.57 |
| 67 | mfc_divcon_3_std.58 |
| 68 | mfc_divcon_5_median.7 |
| 69 | mfc_divcon_5_median.16 |
| 70 | mfc_divcon_5_median.27 |
| 71 | mfc_divcon_5_median.51 |
| 72 | mfc_divcon_5_median.178 |
| 73 | mfc_divcon_5_std.9 |
| 74 | mfc_divcon_5_std.12 |
| 75 | mfc_divcon_5_std.43 |
| 76 | mfc_divcon_5_std.68 |
| 77 | mfc_divcon_5_std.86 |
| 78 | mfc_divcon_5_std.121 |
| 79 | mfc_divcon_5_std.134 |
| 80 | mfc_divcon_7_median.16 |
| 81 | mfc_divcon_7_median.104 |
| 82 | mfc_divcon_7_median.115 |
| 83 | mfc_divcon_7_median.143 |
| 84 | mfc_divcon_7_std.37 |
| 85 | mfc_divcon_7_std.48 |
| 86 | mfc_divcon_7_std.61 |
| 87 | mfc_divcon_7_std.77 |
| 88 | mfc_divcon_7_std.91 |
| 89 | mfc_divcon_7_std.152 |
| 90 | mfc_divcon_7_std.160 |
| 91 | mfc_max.15 |
| 92 | mfc_max.27 |
| 93 | mfc_variance.26 |
| 94 | mfc_variance.35 |
| 95 | mfc_variance.36 |
| 96 | mfc_variance.38 |
| 97 | mfc_variance.39 |
| 98 | mfc_begin.23 |
| 99 | mfcc_delta_divcon_7_mean.6 |
| 100 | mfcc_delta_divcon_7_mean.106 |
| 101 | mfcc_delta_divcon_3_median.2 |
| 102 | mfcc_delta_divcon_3_median.14 |
| 103 | mfcc_delta_divcon_3_std.58 |
| 104 | mfcc_delta_divcon_5_std.47 |
| 105 | mfcc_delta_divcon_5_std.96 |
| 106 | mfcc_delta_divcon_7_median.21 |
| 107 | mfcc_delta_divcon_7_median.125 |
| 108 | mfcc_delta_divcon_7_std.136 |
| 109 | mfcc_delta2_mean.11 |
| 110 | mfcc_delta2_std.17 |
| 111 | mfcc_delta2_divcon_3_mean.20 |
| 112 | mfcc_delta2_divcon_3_mean.28 |
| 113 | mfcc_delta2_divcon_3_mean.29 |
| 114 | mfcc_delta2_divcon_3_mean.119 |
| 115 | mfcc_delta2_divcon_5_mean.20 |
| 116 | mfcc_delta2_divcon_5_mean.26 |
| 117 | mfcc_delta2_divcon_5_mean.132 |
| 118 | mfcc_delta2_divcon_7_mean.166 |
| 119 | mfcc_delta2_divcon_7_mean.229 |
| 120 | mfcc_delta2_divcon_3_median.44 |
| 121 | mfcc_delta2_divcon_3_median.90 |
| 122 | mfcc_delta2_divcon_3_std.37 |
| 123 | mfcc_delta2_divcon_3_std.95 |
| 124 | mfcc_delta2_divcon_5_median.138 |
| 125 | mfcc_delta2_divcon_5_median.149 |
| 126 | mfcc_delta2_divcon_5_std.69 |
| 127 | mfcc_delta2_divcon_5_std.98 |
| 128 | mfcc_delta2_divcon_7_median.13 |
| 129 | mfcc_delta2_divcon_7_median.22 |
| 130 | mfcc_delta2_divcon_7_median.23 |
| 131 | mfcc_delta2_divcon_7_median.28 |
| 132 | mfcc_delta2_divcon_7_median.122 |
| 133 | mfcc_delta2_divcon_7_median.139 |
| 134 | mfcc_delta2_divcon_7_median.151 |
| 135 | mfcc_delta2_divcon_7_median.234 |
| 136 | mfcc_delta2_divcon_7_median.254 |
| 137 | mfcc_delta2_divcon_7_std.65 |
| 138 | mfcc_delta2_divcon_7_std.180 |
| 139 | mfcc_delta2_divcon_7_std.188 |
| 140 | mfcc_delta2_divcon_7_std.201 |

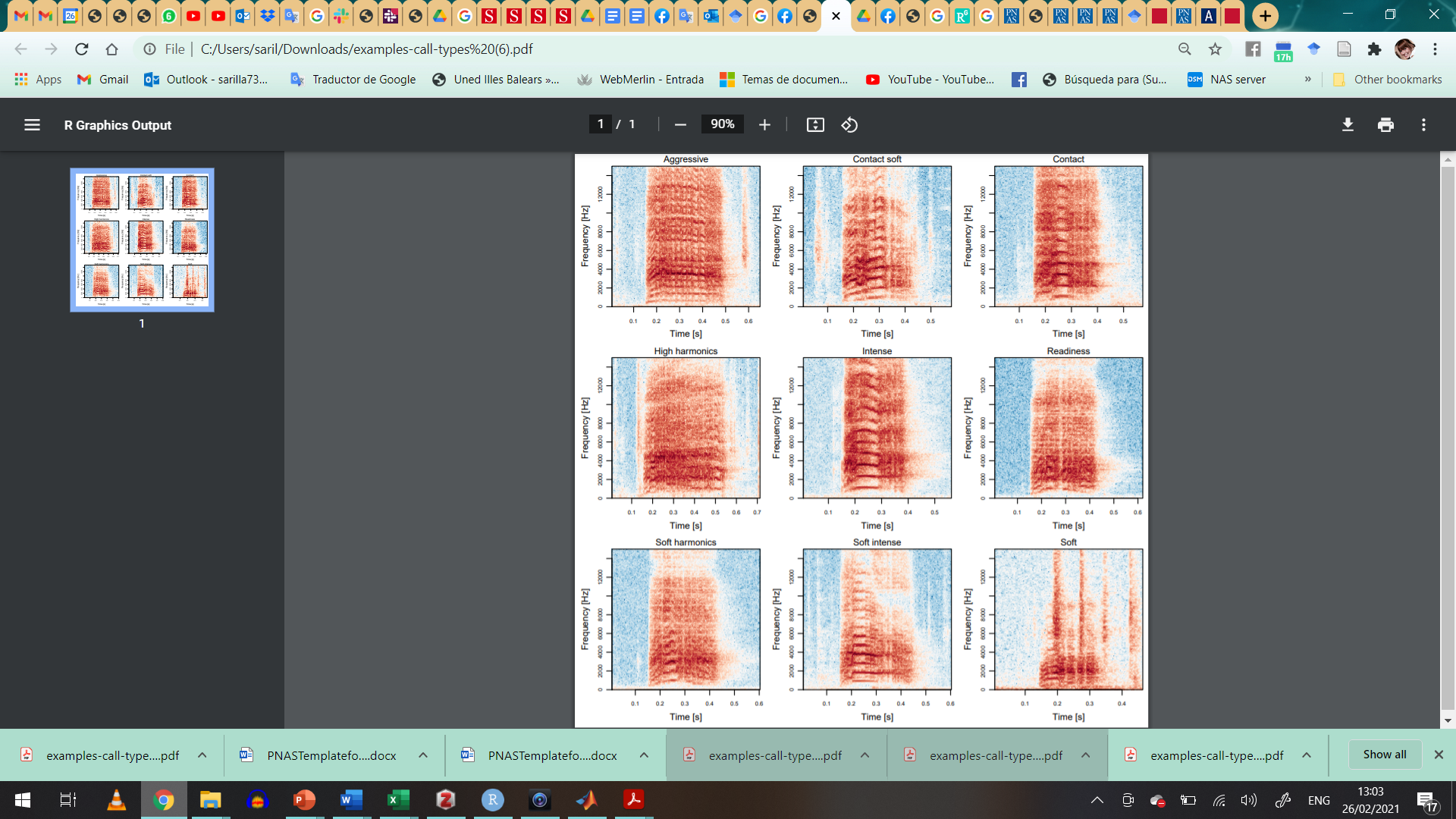

**Figure S1.** Spectrogram for the nine call types shared by all four birds.

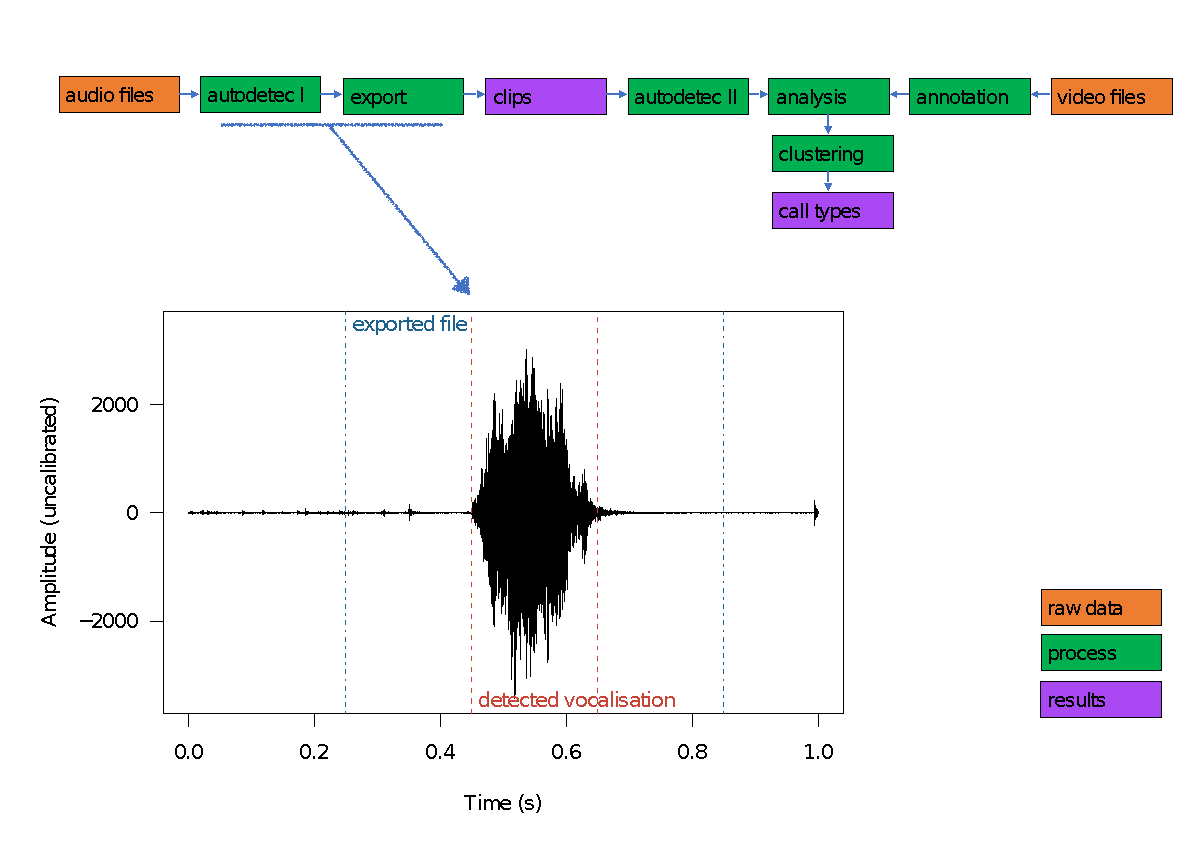

**Figure S2.** Example of processing of raw recordings.

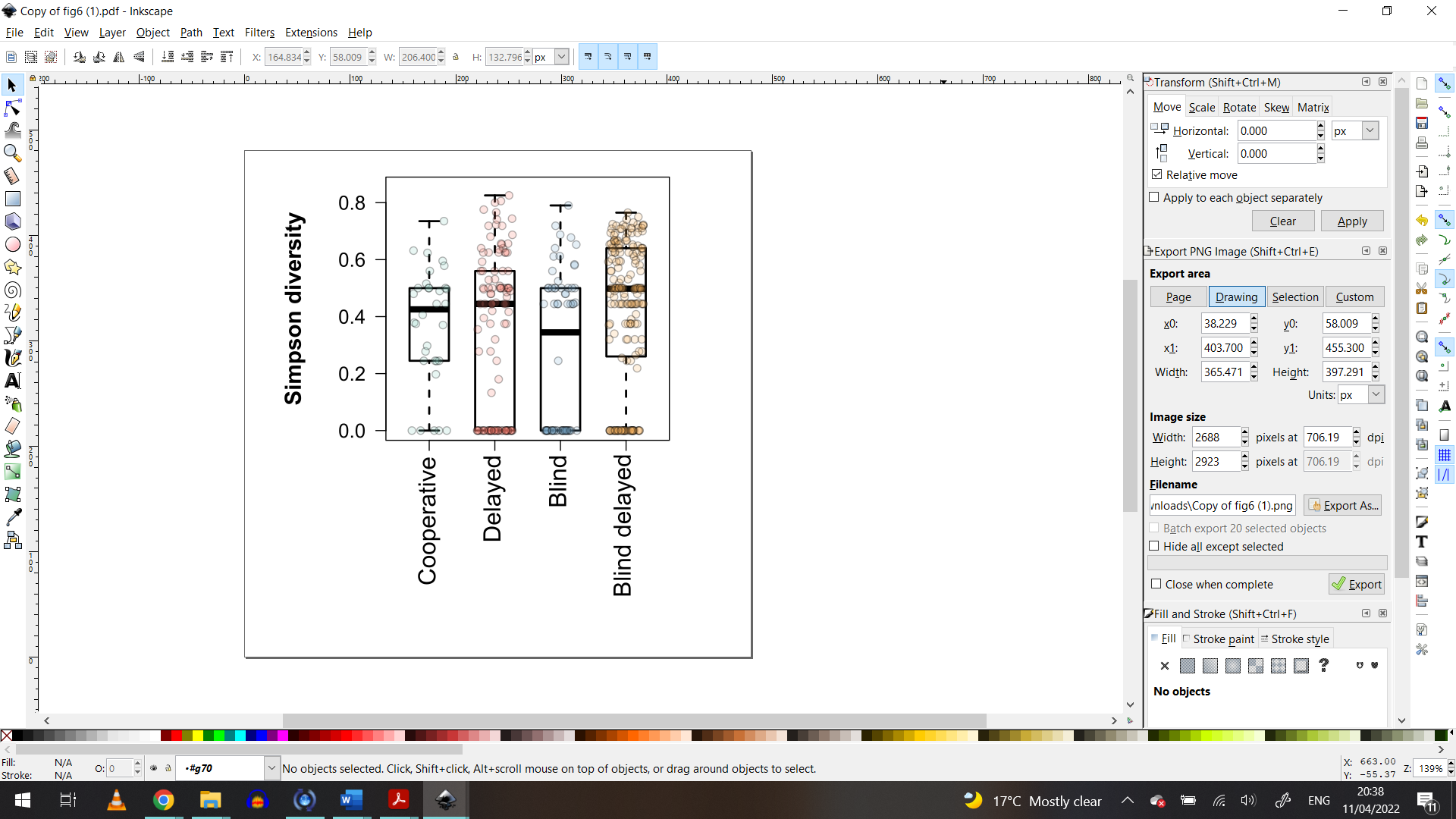

**Figure S3.** Simpson diversity index for the calls during the four experimental conditions. Simpson index was used to measure the diversity taking into account the number of call types present in each condition, as well as the relative abundance of each call type.
